# Unexpected higher convergence of human-great ape enteric viromes in central African forest than in a European zoo: A One Health analysis

**DOI:** 10.1101/2022.07.29.501976

**Authors:** Victor Narat, Maud Salmona, Mamadou Kampo, Thibaut Heyer, Severine Mercier-Delarue, Noémie Ranger, Stephanie Rupp, Philippe Ambata, Richard Njouom, François Simon, Jérôme Le Goff, Tamara Giles-Vernick

## Abstract

Human-animal pathogenic transmissions threaten both human and animal health, and the processes catalyzing zoonotic spillover and spillback are complex. Prior field studies offer partial insight into these processes but overlook animal ecologies and human perceptions and practices facilitating human-animal contact. Conducted in Cameroon and a European zoo, this holistic study elucidates these processes, integrating metagenomic, historical, anthropological and great ape ecological analyses, and real-time evaluation of human-great ape contact types and frequencies. Surprisingly, we find more enteric virome sharing between Cameroonian humans and great apes than in the zoo, a virome convergence between Cameroonian humans and gorillas, and adenovirus and enterovirus taxa as most frequently shared between Cameroonian humans and great apes. In addition to physical contact from hunting, meat handling and fecal exposure, overlapping human cultivation and gorilla pillaging in forest gardens explain these unexpected findings. Our multidisciplinary study identifies environmental co-use as a complementary mechanism for viral sharing.

## Introduction

Pathogenic sharing between human and wild animal populations constitute major threats to human and animal health^1,2^. The virus SARS-CoV-2 illustrates the worst consequences of anthropozoonosis (zoonotic spillover) and the risks associated with zooanthroponosis (spillback): SARS-CoV-2 apparently emerged from bat populations and probably infected other intermediate animal hosts before moving into human populations, causing significant morbidity and mortality; zooanthroponosis may create animal reservoirs that can generate new SARS-CoV2 variants^3–5^. Pathogenic spillback from humans into wild animals, moreover, can hamper conservation of protected species. Such multispecies risks may catalyze new pandemics in the future^6,7^.

The processes leading to anthropozoonoses are complex, driven by genetic proximity between hosts, a pathogen’s adaptative capacity, and human-animal contact, itself catalyzed by anthropogenic changes, including demographic expansion, land use that fragments habitats, economic changes, wild animal trade, hunting and butchering^6,8,9^. Modeling studies help to predict where, when, and who is at greatest risk for emerging anthropozoonoses^10^. Although less studied, zooanthroponoses appear to be facilitated by similar practices and processes^2^. Lacking, however, are fine-grained investigations of potential pathogen sharing between animals and people, and the ecologies, processes, and practices sustaining such sharing^11^.

Existing evidence and analyses only partially illuminate the dynamics of zoonotic spillovers and spillbacks. Identifying potential pathogens at risk of bidirectional sharing remains challenging and expensive^12,13^. Many analyses focus on single pathogens, genera or families^14,15^ and overlook granular evidence of variable human-animal interactions facilitating or mediating against pathogen sharing. Certain human-animal contact investigations elucidate these cross-species interactions, but have been weakened by imprecise definitions of specific contact types and exclusive attention to blood-borne transmission through hunting and butchering, neglecting other transmission modes^16,17^. Finally, investigations integrating host animal ecologies are rare.

This article bridges viral, ecological, and anthropological investigation to produce holistic insight into potential pathogen sharing and the complex interactions facilitating it. It compares the gut virome of humans, gorillas, and chimpanzees residing in two different sites (central African forest and European zoo), detailing their attendant socio-ecological systems^18^ to explain how viral sharing may occur. We focus on human-great ape interactions because of the deep evolutionary relations between these primate species, long histories of shared pathogens (including those originating in different animal species^19^), and frequent interactions between humans and great apes in forest and zoo settings. We analyze the gastrointestinal virome for two reasons: biological collections were noninvasive for protected great ape species, and stools contain important quantities of environmentally-persistent viruses that can facilitate indirect transmission.

We hypothesize that host species and environment will influence the intestinal virome, as other intestinal bacteriome studies have shown^20–24^. We also predict that in both sites, the human virome would more closely resemble that of chimpanzees because of phylogenetic proximity, and because of close daily physical and environmental contact, viral sharing between humans and both great apes would be greater in the zoo than in the Cameroonian forest.

The first and principal site of investigation, located in the southeastern Cameroonian dense rainforest, is home to rural people who derive their livelihoods from farming, gathering, hunting and fishing and who live in close proximity to and share forest and farming spaces with sympatric species of lowland gorillas *(Gorilla gorilla gorilla)* and chimpanzees *(Pan troglodytes troglodytes).* The second site is a European zoo with sympatric chimpanzees *(P.t. verus)* and gorillas *(G.g. gorilla)* and their human zookeepers. This site served as a positive control for human-great ape contact. There, physical and environmental contact is high, continuous, and easily observed because of accessibility, offering conditions that facilitate viral sharing^25^.

The present study is a first comparing across two sites the shared gastrointestinal virome between humans and two nonhuman primate (NHP) species living in close proximity. Our metagenomic sequencing of human and great ape intestinal virome illuminates new, shared, potentially pathogenic viruses^4^. The human virome contains less-explored viral communities that are associated with disease conditions, trigger immune response, or may function as commensals,^26,27^ and few published studies have explored great ape gastrointestinal virome, pathogenic viruses and disease^28–30^. Crucially, our multidisciplinary analysis identifies gorilla and human co-use of small forest gardens as a complementary mechanism for viral sharing, in addition to physical contact through great ape hunting, meat handling and fecal exposure.

We begin with perspectives of people inhabiting the Cameroonian forest concerning their understandings of their history with great apes and socio-cultural perspectives that frame their current relations with gorillas and chimpanzees. We then quantify environmental and physical contacts between humans and great apes. These findings enable an interpretation of our intestinal virome results, in which we document the degree of intestinal virome-sharing between humans and great apes and chimpanzee-gorilla virome differences. Based on this One Health approach, we examine the relative influences of host phylogeny and proximity, ecology, and human perceptions and practices to identify mechanisms for cross-species viral sharing.

## Results

### Humans and great apes in Cameroon: a lengthy, shared history

Our qualitative evidence on past and contemporary human-great ape relations reveals that Southeastern Cameroonians believed that they shared a lengthy history with great apes. Several mythical tales *(likano),* still recounted during evening festivities, portray a distant past in which people, chimpanzees, gorillas, and gods once lived together in settlements (Likano sessions, 12.07.2015, 08.07.2015). These tales indicate that cohabitation was ruptured when chimpanzees and gorillas committed social transgressions, resulting in their ejection from human society.

This deep history of cohabitation and interaction frames local assumptions that people and great apes share specific capacities. For our informants, great ape displays of certain emotions through their protection of other troupe members, mutual grooming, and sorrow reflected their commonalities with human capacities for love, emotional expression, and mutual protection. Great apes, however, displayed superior strength to humans, and interviews with local healers reveal that this strength rendered certain body parts (tibia or vertebrae of chimpanzees, bone marrow from gorillas) useful in remedies for human weaknesses or ailments (Interviews 21.05.2016, 20.01.2016). The frequency of these uses is unknown.

These long-term histories framed current human-great ape interactions in other ways. Mythical tales recounting past human-great ape cohabitation, ruptured by great apes’ exile, was a narrative structure mirrored in recent oral histories. Oral accounts underscored physical and emotional distancing between people and great apes that accelerated over the last 50 years (Interviews 15.07.2015, 29.05.2017, 30.05.2017, 02.06.2017, 30.04.2016). As one aging hunter observed,

> Before...gorillas slept next to the village. Now, they are there, but very rarely next to the village. They are afraid and very shrewd. Even the chimpanzees, there’s a group between this and the next village...just a few meters from the road. But they don’t cry out, they are prudent. It’s rare to hear them, even though they are there (Interview 15.07.2015).

For some, this distancing resulted from intensified hunting since the 1970s with high-powered rifles (Interviews 13.05.2016, 14.05.2016, 02.06.2017). Perceptions of the changing sizes of gorilla and chimpanzee populations are mixed: some informants contend that great ape populations have remained stable, and others argue that they have declined (Interviews 29.05.2017, 30.05.2017, 21.01.2016).

### Differential human contacts with chimpanzees and gorillas in Cameroon

Although Southern Cameroonians generally assumed that they shared a deep history with great apes, and over time were distanced from them, they did not recognize “great apes” as an NHP category. They distinguished chimpanzees *(waké)* from gorillas *(ko),* engaging with chimpanzees differently and less frequently than with gorillas.

Our data show that southeastern Cameroonians perceived chimpanzees as more intelligent than gorillas, capable of learning and displaying behaviors that approximate, but do not replicate, human behavior. Chimpanzees were more elusive than gorillas, living close to human settlements but remaining silent and invisible. Some specialist hunters also claimed that it was easier to kill a gorilla than a chimpanzee, contending that this relative ease partly resulted from gorillas’ large size and practice of moving across the forest floor, rendering them more detectable than chimpanzees, who frequently traversed tree canopies (Interviews 31.C5.2C17, 18.C1.2C16). They also reported that emotionally, it was more difficult to kill a chimpanzee than a gorilla, because the former more closely resembled humans. As one former hunter confided, “You need a strong heart to kill a chimpanzee.”

Southeastern Cameroonians reported that gorillas were less discerning, clumsier, more destructive, more frequently encountered than chimpanzees, and more unpredictably aggressive (Interviews 18.01.2016, 19.01.2016, 20.01.2016, 21.01.2016). Reportedly unable to distinguish ripe from unripe foods, gorillas more often laid waste to forest gardens, whereas chimpanzees purportedly raided only ripe foods from these gardens. Among people responding to a questionnaire conducted in four villages, 94% (424/449) experienced crop raiding by all NHPs, including gorillas and chimpanzees. Gorillas were more involved in the last event of crop raiding (50%, 225/449), contrary to chimpanzees (2%, 7/449). Moreover, 82% of respondents (366/449) reported that gorillas more often damaged gardens, and just 2% (7/449) cited chimpanzees as more frequent pillagers.

Quantitative data on human physical and environmental contacts, collected in real time, also illuminates different human interactions with gorillas and chimpanzees. The mean frequency of human physical and environmental contact with gorillas was higher, but not significantly, than with chimpanzees; direct contact (seen alive, heard) did not differ between these great apes (Table 1). Based on questionnaires, physical contacts were more frequent (except for hunting) with gorillas than chimpanzees.

**Table 1:** Mean (%) contact frequencies (SD) according to great ape species and type of contact in Cameroon. Physical contact frequencies are from Narat et al. 2018. *** indicates a p-value <0.001 from the Mann-Whitney statistical tests.

|  |  | Longitudinal survey<br>N=18, data self-collected daily,<br>10 months |  | Questionnaire<br>N=449 |  |
| --- | --- | --- | --- | --- | --- |
| Contact category | Type of contact | *Pan t. troglodytes* | *Gorilla g. gorilla* | *Pan t. troglodytes* | *Gorilla g. gorilla* |
| Environmental contact | Seen feces | 2.3 (5.3) | 3.5 (5.3) | Not addressed in questionnaires |  |
|  | Seen food remains | 2.5 (5.3) | 4.0 (5.6) |  |  |
|  | Seen nest | 1.9 (3.5) | 1.5 (2.3) |  |  |
|  | Seen footprints | 2.2 (5.2) | 3.8 (5.3) |  |  |
| Direct contact | Seen alive | 1.8 (5.1) | 1.8 (4.1) | 14.2 (31.4) | 9.1 (22.3) |
|  | Heard | 3.2 (5.3) | 2.6 (4.2) |  |  |
| Physical contact | Hunt | 0 (0, 0-0) | 0 (0, 0-0) | 0.08 (0.7) | 0.07 (0.3) |
|  | Butcher | 0.1 (0.3) | 0.2 (0.3) | 0.7 (2.2)*** | 1.6 (7.3)*** |
|  | Cook | 0.1 (0.4) | 0.2 (0.6) | 0.6 (2.1)*** | 1.0 (2.7)*** |
|  | Consume | 0.2 (0.5) | 1.1 (2.5) | 0.7 (2.2)*** | 1.7 (7.3)*** |
|  | Buy/Sell | 0.5 (1.2) | 1.1 (2.6) | 0.4 (1.6)*** | 1.3 (7.1)*** |

### Human-great ape contact at the European zoo

Zoo gorillas and chimpanzees lived separately but shared the same environment. Both species had an indoor enclosure with outdoor access on an island with similar vegetation. Islands were surrounded by channels through which water continually circulated. Our observations showed that zookeepers and great apes had notable daily environmental and physical contact. Although we did not quantify their contact frequency, we observed daily zookeepers in close proximity (<2m) to gorillas and chimpanzees, especially through the bars of cages. Zookeepers handled food during preparation to feed to chimpanzees and gorillas. Moreover, before the COVID-19 pandemic, they generally did not wear masks and gloves when entering the cages each day to remove straw bedding and feces, and when hosing cages with hot water every 3 to 5 days.

We observed occasional physical contact between zookeepers and chimpanzees (playing, affectionate scratching, grooming), whereas no physical contact occurred between zookeepers and gorillas. Additionally, zookeepers reported that chimpanzees fell ill more often than gorillas, especially with respiratory infections during winter months.

### Fecal sample characterization and comparison

We focused on vertebrate viruses because of their importance for viral transmission and emergence. Among vertebrate viruses, 13 families, 26 genus and 61 species were identified. The most frequent viral families observed in decreasing order were Adenoviridae (53 samples), Picobirnaviridae (30 samples) and Picornaviridae (29 samples). The distribution of viral vertebrates reads per family for each sample tested is shown in Figure 1. The distribution of viral reads across their natural host categories is depicted in Supplementary Figure 1.

**Figure 1:**
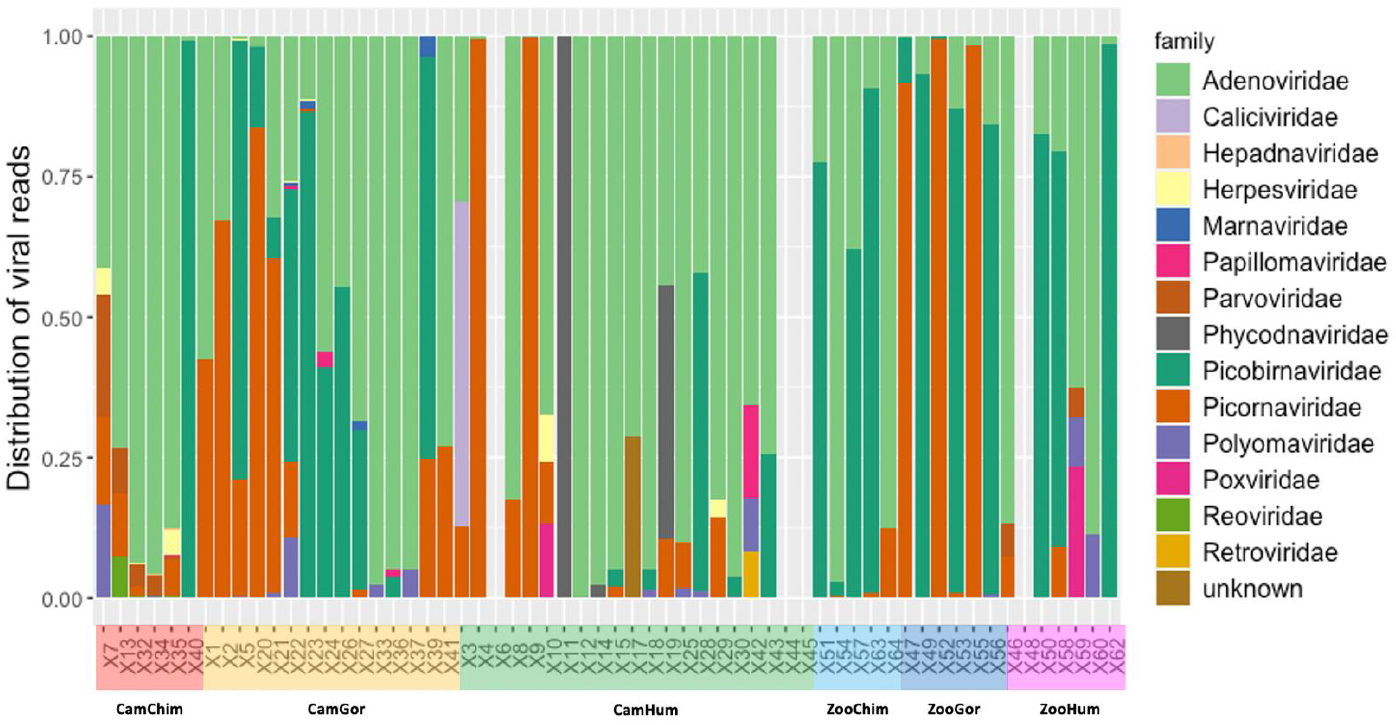
Distribution of viral vertebrate reads per viral family. Each group is defined by its habitation site (Cam=Cameroon, Zoo=European Zoo) and animal species (Chimp=Chimpanzee, Gor=Gorilla, Hum=Human)

The number of samples positive for each virus family among the ten most prevalent viral families is detailed in Table 2. Adenoviridae, Picobirnaviridae and Picornaviridae were identified in all human, gorilla and chimpanzee groups and Polymaviridae in all but zoo chimpanzees. Herpesviridae, Papillomaviridae and Parvoviridae were identified only in stool samples of apes and humans in Cameroon.

**Table 2:** Number (%) of samples found positive for one of the 10 most prevalent vertebrate viral families by group. Each group is defined by its habitation site (Cam=Cameroon, Zoo=European Zoo) and animal species (Chimp=Chimpanzee, Gor=Gorilla, Hum=Human).

|  | CamChimp<br>n=6 | CamGor<br>n=15 | CamHum<br>n=23 | ZooChimp<br>n=5 | ZooGor<br>n=6 | ZooHum<br>n=7 |
| --- | --- | --- | --- | --- | --- | --- |
| **Adenoviridae** | 6 (100%) | 14 (93%) | 17 (74%) | 5 (100%) | 5 (83%) | 6 (86%) |
| **Caliciviridae** |  |  | 1 (4%) |  |  |  |
| **Hepadnaviridae** | 1 (17%) |  |  |  |  |  |
| **Herpesviridae** | 4 (67%) | 3 (20%) | 2 (9%) |  |  |  |
| **Papillomaviridae** | 1 (17%) | 2 (13%) | 1 (4%) |  |  |  |
| **Parvoviridae** | 5 (20%) | 1 (7%) | 2 (9%) |  |  |  |
| **Picobirnaviridae** | 2 (33%) | 10 (67%) | 6 (26%) | 4 (80%) | 5 (83%) | 3 (43%) |
| **Picornaviridae** | 4 (67%) | 9 (60%) | 10 (43%) | 2 (40%) | 3 (50%) | 1 (14%) |
| **Polyomaviridae** | 2 (33%) | 5 (33%) | 5 (22%) |  | 1 (17%) | 2 (29%) |
| **Poxviridae** |  |  | 1 (4%) |  |  | 1 (14%) |

### Viral diversity and composition

Global virus richness was significantly higher in Cameroon chimpanzees than among Cameroonian humans or zookeepers (Observed and Chao index), although no difference between these same groups was observed with Shannon and Simpson indexes (Supplementary Figure 2a). When we focused only on vertebrate viruses, no significant difference was observed between the six different groups, despite a tendency for higher diversity in Cameroon chimpanzees (Supplementary Figure 2b).

The comparisons of virome composition using a PERMANOVA with Bray Curtis dissimilarity indices and weighted unifrac showed that the virome differed significantly across different groups (p values < 0.001). An unsupervised analysis with a PCA among all samples showed that the global virome composition differed between host species, with a closer similarity between zoo chimpanzees and gorillas (Figure 2a and 2b). The network projection of virome similarity confirmed the proximity between viromes of zoo great apes. Viromes of zoo great apes resembled that of Cameroonian chimpanzees, which in turn was close to that of Cameroonian gorillas. Despite the distinct environments in which stools were collected, the Cameroonian and zoo human viromes closely resembled one another. Surprisingly, the network analyses revealed greater virome resemblance between Cameroonian humans and Cameroonian gorillas than between Cameroonian gorillas and other great apes in the Cameroon forest or the zoo (Figure 2c).

**Figure 2:**
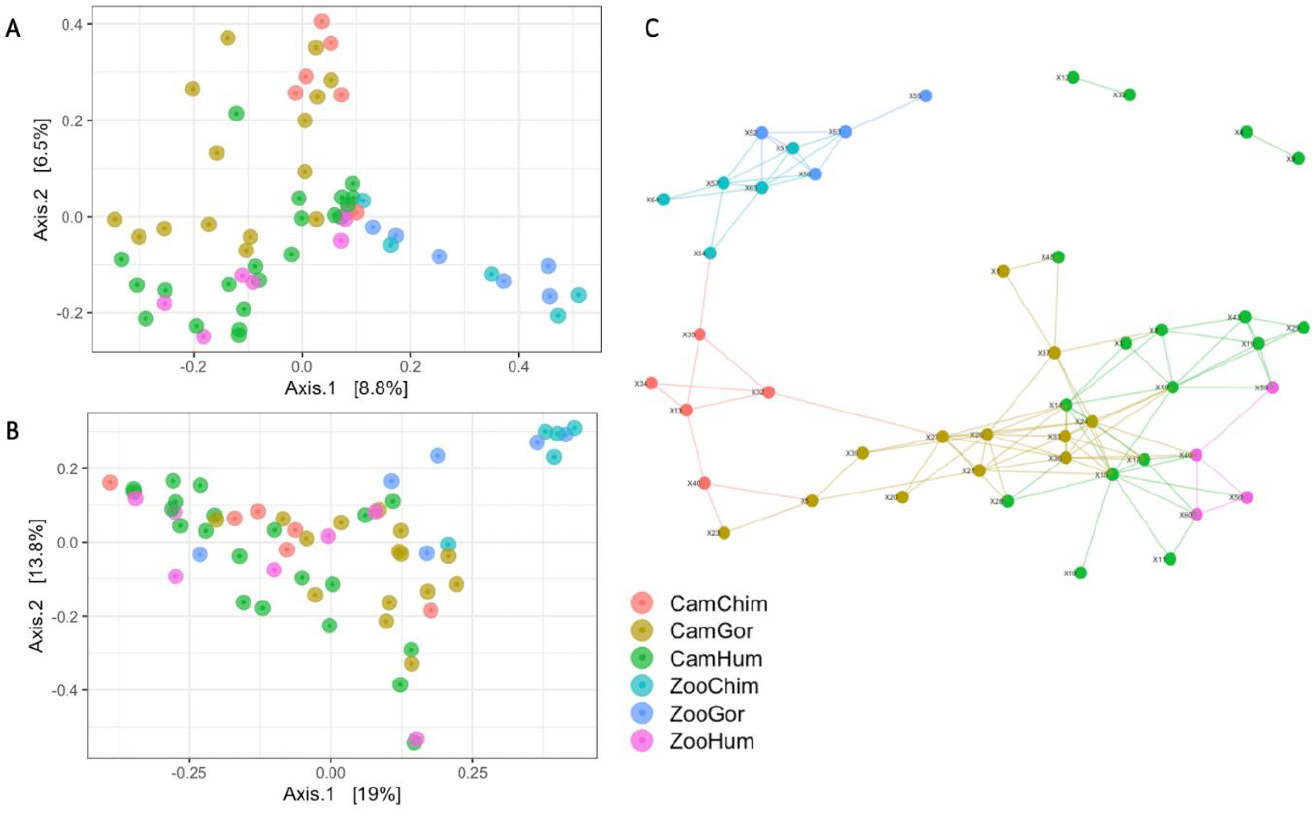
PcoA analysis based on Bray-Curtis distances (A) and on weighted Unifrac distances (B). (C) Network plot based on Bray-Curtis distances showing similarities among all sample virome profiles. Only edges connecting individuals (i.e., nodes) with > 90% similarity in their virome are shown. Each group is defined by its habitation site (Cam=Cameroon, Zoo=European Zoo) and animal species (Chimp=Chimpanzee, Gor=Gorilla, Hum=Human)

### Viral sharing

We then investigated whether within a shared environment, great apes and humans might harbor viruses with same lowest-common-ancestor (LCA) taxa identified. In Cameroon, a total of 15 vertebrate viral LCA taxa identified in human stool samples were also found in those of chimpanzees or gorillas: 13 viral LCA taxa in gorilla and human stools, 8 in chimpanzee and human stools, and 5 shared by all 3 groups (Figure 3a). Among these vertebrate viral taxa, the most represented genera were Mastadenovirus (n=5), Picobirnavirus (n=3), and Enterovirus (n=3). In the European zoo, 8 viral LCA taxa were shared between humans and apes, with a greater representation of the Picobirnavirus genus (n=5) (Figure 3b).

**Figure 3:**
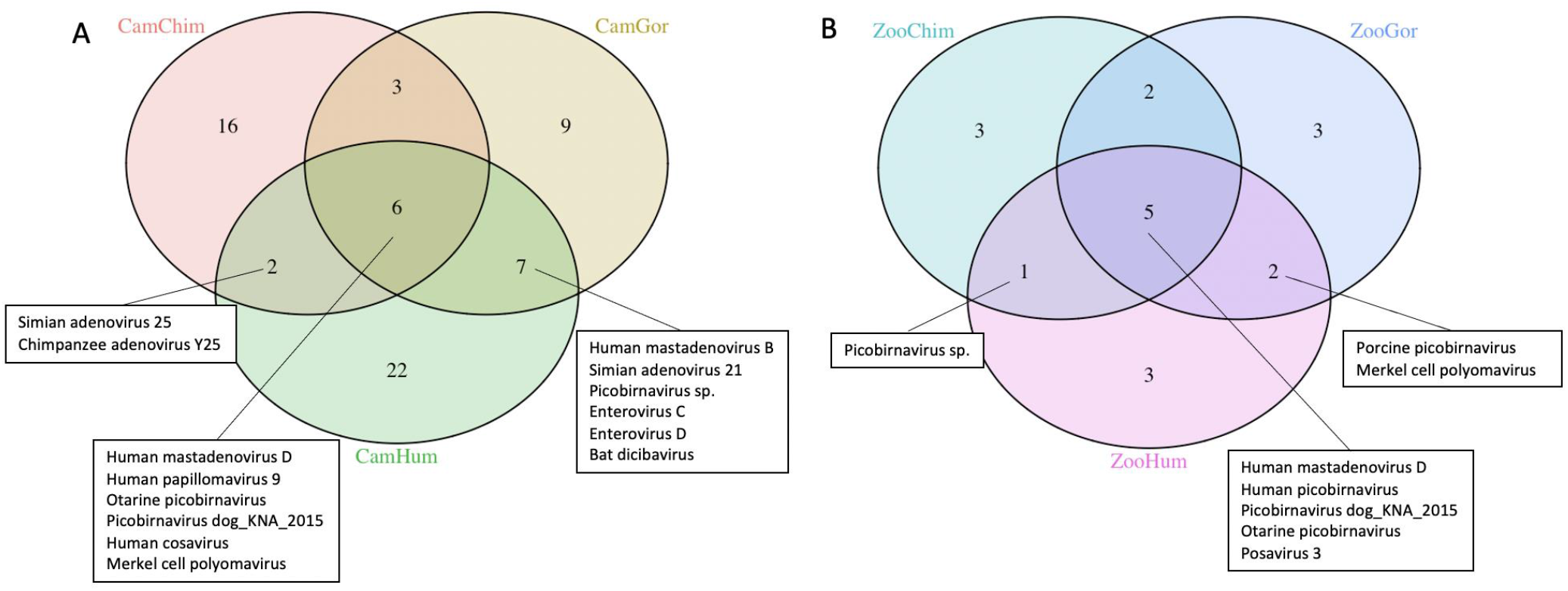
Venn Diagrams of vertebrates viral lowest-common-ancestor (LCA) taxa identified in human and great ape stools in Cameroon (A) and European zoo (B). Each group is defined by its habitation site (Cam=Cameroon, Zoo=European Zoo) and animal species (Chimp=Chimpanzee, Gor=Gorilla, Hum=Human)

We analyzed the Mastadenovirus and Enterovirus genera, the vertebrate viruses that great apes and humans shared. Concerning the Mastadenovirus genus, we detected adenovirus species D among humans and great apes in Cameroon and in the European zoo. We found species E in Cameroonian humans, gorillas and chimpanzees, and species B in Cameroonian humans and gorillas. Although we detected Adenovirus species B and E in zoo great apes, we did not find them among zookeepers. Enterovirus C and D species were detected only in Cameroonian humans and gorillas.

To evaluate the genetic proximity of the viral sequences, reads belonging to Mastadenovirus and Enterovirus genera were assembled to create contigs. Because the size of the contigs did not permit building of phylogenetic trees to compare different viruses, a network analysis on cytoscape was performed to assess genetic similarities between shared viruses. The sequence similarity network confirmed the detection of adenovirus species B, D and E (Figure 4a). The network showed close genetic distances between some adenovirus D collected among humans, chimpanzees and gorillas (Figure 4a), and between an enterovirus C species collected in one Cameroonian human and those collected from Cameroonian gorillas and chimpanzees (Figure 4b).

**Figure 4:**
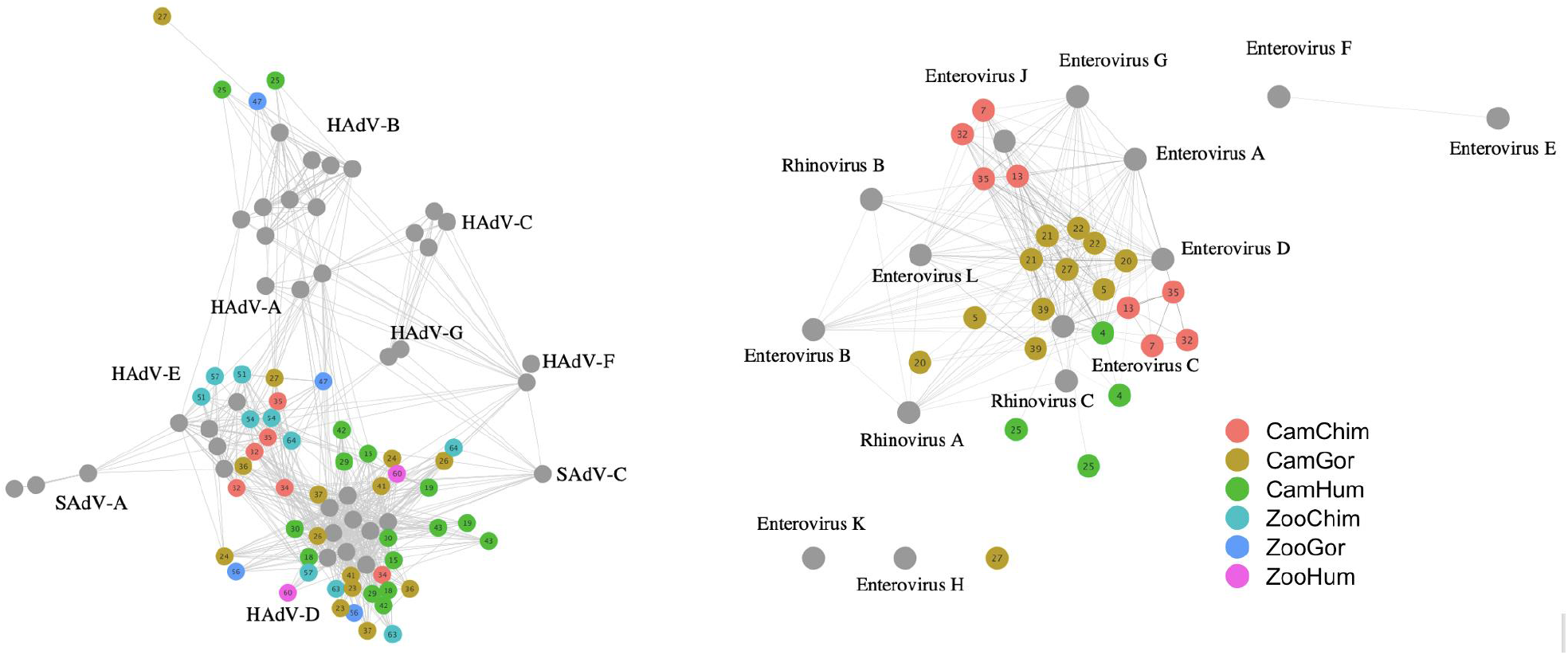
Sequence similarity network representation of Mastadenovirus (a) and Picornavirus (b) contigs. Each node represents an individual contig or whole genome reference sequences. Edges are defined based on the Blast Bit-score across individual samples. Each contigs is colored according to the group to which it belongs. Grey nodes represent reference sequences. Each group is defined by its habitation site (Cam=Cameroon, Zoo=European Zoo) and animal species (Chimp=Chimpanzee, Gor=Gorilla, Hum=Human)

## DISCUSSION

To shed light on shared human-great ape viromes and possible mechanisms of bidirectional viral sharing, this study brings together metagenomics analyses, oral histories, anthropological observation, great ape ecological analyses, and real-time evaluation of contact types and frequencies between people and great apes living in the African equatorial rain forest. It compares these results to a positive control in a European zoo, where we easily observed zookeeper-gorilla and zookeeper-chimpanzee interactions. We had two unexpected findings: the convergence of intestinal virome between Cameroonian humans and gorillas, and the higher proportion of human-great ape sharing in Cameroon. We expected that humans would share more of their intestinal virome with chimpanzees because of their phylogenetic proximity, and that viral sharing would be greater in the zoo, our positive control, because human zookeepers had closer, more frequent contact with zoo great apes. Additionally, certain adenovirus and enterovirus species were the most frequently shared viruses between Cameroonian great apes and humans. Our multidisciplinary analyses explain these biological findings in terms of southern Cameroonians’ distinct perceptions and practices toward gorillas and chimpanzees, and gorilla and chimpanzee relative densities and behaviors.

The following explains each of our analyses around human-great ape interactions and risks associated with two viral genera, Mastadenovirus and Enterovirus, shared by Cameroonian humans and great apes. We then integrate these analyses to identify complementary mechanisms of human-great ape viral sharing in the central African forest: physical contact through hunting, meat handling, and fecal exposure; and environmental contact concentrated in small forest gardens, frequented by humans and great apes, especially gorillas.

### A historical framework for human-great ape interactions

Although such evidence is not habitually integrated into metagenomics studies, our qualitative findings offer a historical framework that describes and still shapes human-great ape relations. Southeastern Cameroonians maintain that for millennia, they have interacted and shared forest spaces and foods with great apes; historical linguistic and archaeological investigations support this claim^31,32^. People now interact less with great apes than in previous generations, but gorillas and chimpanzees still spark avid interest and ambivalence among southeastern Cameroonians. The compelling nature of great apes appears to result from their “charismatic” features; their behaviors lend themselves to human observation and elicit strong emotional responses among observers^33^.

Southeastern Cameroonian mythical tales and historical recollections make no reference to biological evolution. Consistent with historical, anthropological interpretations of mythical tales, southeastern Cameroonians appear to have distilled from their accumulated interactions with great apes their reflections about how humans, gorillas and chimpanzees resemble and differ from one another. These tales comment on an increasing emotional and physical distance between people and great apes, although there is little agreement on how it happened^34–36^. In the absence of long-term great ape population surveys, we cannot conclude that these NHP populations have been depleted. Certain people hunt great apes in this forest, whereas others do not^37^.

Our anthropological research shows that southeastern Cameroonians distinguish chimpanzees from gorillas, their perceptions shaping their practices around these great apes. Chimpanzee behavior, they concur, is closer than gorilla behavior to that of humans. Hence, southeastern Cameroonians avoid engaging with chimpanzees and are more reluctant to kill and butcher them, a tendency reinforced by chimpanzees’ avoidance of humans. Gorillas, by contrast, are reportedly more numerous. Their “less human” behavior apparently makes it easier for people to kill and butcher them. Human perceptions of differences between gorillas and chimpanzees seem to yield more frequent human physical and spatial contacts with gorillas than chimpanzees.

### More frequent human-gorilla contact than human-chimpanzee contact in Cameroon

These qualitative findings align well with our quantitative analyses of human-great ape contact frequency. Our participatory longitudinal study found elevated, but not significant, human-gorilla spatial contact compared to that of humans and chimpanzees. Our questionnaire dataset, however, revealed significantly more frequent physical contact with gorillas. Most studies assessing human-great ape contact do not evaluate contact frequency^38–40^, an important but overlooked factor in understanding pathogenic transmission risks. Our previously published study evaluated quantitative and anthropological data concerning human-great ape contact and different contact types with diverse NHP species^41^. Here, integrating quantitative contact analyses and qualitative evidence about such contacts permits insight into how southeastern Cameroonians may share specific viruses with great apes and closer virome resemblance to that of gorillas.

Our Cameroon investigation also describes the context and locations of these frequent human-great ape physical and environmental contacts. Our multi-village questionnaire showed that great apes were frequently involved in garden crop raiding, and gorillas more so than chimpanzees. Several primate ecological and behavioral factors may explain greater physical and environmental contact with gorillas. First, gorillas appear to be more abundant in the Cameroonian forest, leading to a higher probability of gorillas entering and raiding gardens may explain this phenomenon. Our previous work showed that signs of gorilla presence were 10 times higher than for chimpanzees in this forest^41^, in accordance with other studies finding that gorillas are more abundant or in similar densities to sympatric chimpanzees in the Republic of Congo, DR of Congo, Gabon, and Cameroon^42–44^.

In addition to different gorilla and chimpanzee relative densities, our southeastern Cameroonian informants also reported that gorillas crop raided more frequently and destructively. For many animals, including gorillas, these gardens were concentrated sites of food raiding and defecation. Gorillas and other nonhuman primates defecated freely in or around gardens. Gardens, frequently distant from villages where people constructed latrines, could also be sites of human defecation during their daily visits. In both cases, defecation could deposit environmentally persistent microbes.

Gorillas and chimpanzees may behave differently in forest gardens, leading gorillas to raid more often than chimpanzees. Most existing literature NHP crop raiding, however, focuses on chimpanzees in east and west Africa^45–48^. Just two studies address mountain gorilla *(Gorilla beringei)* crop raiding^45,48^. Gorilla and chimpanzee behavioral differences in socio-ecological systems require further investigation^49^. Nonetheless, small gardens may be focused “sharing platforms” for environmentally persistent microbes and their exchanges between humans and gorillas.

### Influence of host species and environment on intestinal virome

Our results indicate that each species in its own environment harbored a specific virome composition, as expected. As with gut bacterial microbiota, phylogeny is a strong driver of species-specific enteric virome^20,22,50^. Our findings corroborate a previous investigation of the evolutionary and ecological origins of gut bacteriophage communities (phageome), demonstrating that the phageome structure and dynamics were influenced by superhost phylogeny and environment^51^.

Network and dissimilarity analyses revealed that human enteric viromes from Cameroon forest and the zoo were more similar despite habitation in different biotopes, whereas virome composition for great apes appeared to be shaped more significantly by ecology than by species. Additionally, the zoo environment seems to have exercised a greater influence on great ape virome than the forest did for Cameroon great apes. These results corroborate those reported by Moeller and colleagues^52^, in which gut microbiota of sympatric chimpanzees and gorillas bore greater resemblance to one another than gut microbiota of either allopatric bonobos and eastern lowland gorillas or allopatric chimpanzees from Tanzania and eastern lowland gorillas. Hence, our findings on human and great ape gut viromes reveal similar patterns to comparative gut microbiomes. Although phylogeny seems to exercise a greater impact on human gut virome and microbiome, environment apparently has a stronger influence on great ape viromes.

### Targeted assessment of bidirectional viral sharing risks

Among vertebrate viruses known to cause disease in human beings, we identified Mastadenoviruses and Enteroviruses as the major viral genera that humans and great apes share in Cameroon. Both genera can be transmitted through physical contact and can remain infectious in the environment for several days.

Mastadenoviruses are associated with many diseases, including mild and severe respiratory infections, gastro-enteritis, encephalitis, cystitis, keratoconjunctivitis and hepatitis. Recently, severe hepatitis cases of unknown etiology among young children have been reported. Because an adenovirus has been detected frequently in these patients’ feces or blood, investigators have hypothesized that the etiology of the severe hepatitis is an adenovirus, although investigations are ongoing^53^.

Adenovirus is a non-enveloped virus with a double-stranded DNA genome. Primate adenoviruses belong to the genus Mastadenovirus, which includes seven species (A to G) and more than 100 different types. Our analyses found adenovirus species D to be most frequently shared between humans and apes. Adenovirus species D displays frequent recombination events between different types. This tendency to recombine enables the emergence of new types that could escape from previously acquired anti-adenovirus host immunity and potentially trigger disease outbreaks in humans or great apes. Although adenoviruses are considered highly specific to hosts because of the genome’s DNA structure, great ape-human transmission of adenovirus has been shown. A novel adenovirus (TMAdV, titi monkey adenovirus) discovered at the California National Primate Research Center caused a deadly outbreak in a closed colony of New World monkeys (titi monkeys; *Callicebus cupreus)* and infected humans in close contact^54^. Human-to-human transmission of TMAdV was also documented. Other studies have detected specific antibodies of baboon and chimpanzee adenoviruses among caregivers and zookeepers, confirming the capacity of NHP adenoviruses to infect humans^55,56^. Humans may also be capable of transmitting adenoviruses to great apes. Adenoviruses detected among NHPs elsewhere in Cameroon revealed sequences closely related to human adenoviruses^57^. Although no large adenovirus epidemics have been reported, these observations and our findings suggest an elevated risk of cross-species outbreaks.

Recently, machine learning analyzing viral genomes has been used to predict viral zoonotic risk. One study found significantly elevated predicted zoonotic risk in viruses from NHPs and identified adenoviruses among those viruses correlated with the probability of human infection, confirming prior investigations showing that NHP adenoviruses and retroviruses, bat rhabdoviruses, and rodent picornaviruses were more likely to be zoonotic^58,59^.

The second most frequent viral genus shared between humans and great apes was enterovirus, known to cause mild to severe disease in humans. Enteroviruses are positive-sense single-stranded RNA viruses with high mutation rates, frequent recombination events, and a potential for newly emerging types. Based on detection of specific antibodies or targeted PCR and specific viral protein sequencing, a few studies provide some evidence for anthropozoonosis and zooanthroponosis^60^. Although enterovirus origins are difficult to ascertain because most have been described in humans, some studies have detected human enteroviruses in zoo-housed and free-ranging NHPs, with variable frequency depending on housing conditions or the degree of cohabitation in urban areas. In urban Bangladesh where NHPs and humans share the same environment, 100% of enteroviruses detected in NHPs were also known to circulate among the human population^61^. In contrast, in a Bangladesh zoo, most Picornaviruses detected in NHPs (53/64; 83%) were simian viruses, with only 8 (12.5%) detected in humans^62^. These results corroborate our findings, suggesting that environmental sharing without specific hygiene control measures enables cross-species transmission of enteroviruses. Another study investigating enterovirus genetic diversity in 615 stool samples collected between 2006 and 2008 from zoo and free-ranging NHPs in Cameroon, the rate of enterovirus detection was 20.2% among zoo NHPs and 3.5% in free-ranging NHPs. These viruses belong to virus types that circulate among humans in 94% of zoo NHP and 55% of free-ranging NHP positive samples^25^. The zoo NHP habitat (large enclosures), where frequent interactions between NHPs and employees and the public, was suspected as a key site facilitating viral transmission.

### Humans, gorillas, intestinal virome, and targeted viruses: convergences in Cameroon

Contrary to our prediction, viromes of Cameroon forest inhabitants and Cameroon gorillas more closely resembled one another than those of zookeepers and zoo gorillas and chimpanzees. The proportion of viral taxa shared between gorillas and humans was four times higher in Cameroon than in the European zoo, and among Cameroon humans and great apes, human-gorilla viral sharing was higher than for human-chimpanzee sharing.

Our historical-anthropological, ecological, and contact analyses suggest two complementary mechanisms to explain viral sharing: physical contact from hunting, meat handling and fecal exposure, and environmental contact through co-use of and fecal exposure in small forest gardens. Support for these mechanisms comes from multiple findings. Perceiving chimpanzees to resemble humans more closely than gorillas, Cameroonian participants avoided chimpanzees and had more frequent physical contact with gorillas through hunting and meat handling, although such activities were not highly frequent. Gorillas appeared more abundant and more active in crop-raiding than chimpanzees in gardens. Small forest gardens thus constituted focused sites of human-gorilla overlap and environmental contact, where people and gorillas could leave behind environmentally persistent enteric viruses.

The positive control for human-great ape contact and viral sharing – the European zoo – generated unexpected results. Zookeepers and zoo great apes had daily contact and close spatial proximity in a relatively small site, but viral sharing was lower in the zoo than in Cameroon, possibly limited by zookeepers’ occasional handwashing, but also by the above-mentioned mechanisms, and notably environmental contact in Cameroonian gardens.

Targeted viral discovery results are consistent with these two sharing mechanisms. Physical and environmental contact can facilitate sharing of Mastadenoviruses and Enteroviruses and could potentially lead to pathogenic spillover or spillback. An increased frequency of physical and environmental contacts between humans and great apes could facilitate the emergence of a novel viral disease in human or NHPs. Social sciences evaluation of human-NHP contact intensity and the introduction of viral surveillance programs where humans and NHPs are in close engagement would be essential for pandemic prevention.

Finally, our investigation reveals the explanatory richness of multidisciplinary investigations of cross-species pathogen sharing. Although limited to correlations, our anthropological-historical and ecological analyses and our granular study of contact type and frequency were essential for explaining the possible routes for viral sharing, illuminated by our metagenomics analyses and targeted viral discovery.

### Limitations of the study

Our study has several limitations. First, we collected stool samples over one to two weeks, depending on the site and did not repeat collections. This collection strategy may have influenced our metagenomics analyses. We collected our Cameroon samples during the dry season. Seasonal availability of foods may influence virome composition, and in turn, the similarities and differences observed^63^.

Samples were limited in number, primarily because collecting great ape stools in forest settings is challenging. Our samples are, however, numerically sufficient to offer insight into shared gut viromes. We did not collect other NHP stool samples, notably among those with whom Cameroonian people have high-frequency physical or other contact; this investigation focused on great apes because of their greater phylogenetic proximity with humans. Additional sampling would be important for understanding virome sharing between Old World monkeys and human beings in this forest.

Our study investigated the viromes from stool samples and cannot shed light on all potential human-great ape viral transmissions, notably blood-borne or respiratory viruses. Stool samples may, however, include viruses with a non-digestive tropism. Certain respiratory viruses can be detected in stool samples, such as naked viruses (picornavirus, adenovirus) or some enveloped viruses (such coronaviruses, influenza viruses) but usually with lower viral loads than in the respiratory tract^64^. We found no enveloped respiratory viruses in great apes or humans.

## CONCLUSION

In recent decades, research into pathogenic anthropozoonosis and zooanthroponosis has acquired crucial importance. The COVID-19 pandemic has heightened that importance, but also revealed the complex, potentially unexpected spillover and spillback pathways that a virus can take. The present study undertook a multidisciplinary One Health approach to examine shared virome and bidirectional viral sharing between humans and great apes and to illuminate associated processes, practices, and behaviors that may facilitate it. This multidisciplinarity was essential in interpreting our unexpected biological results. More human-great ape sharing in Cameroon than in our control (the European zoo), a surprising convergence of human and gorilla virome in Cameroon, and apparent sharing of adenovirus and enterovirus taxa can only be understood by putting into dialogue metagenomics, historical, anthropological, and ecological analyses in southeastern Cameroon. These analyses point to lengthy human-great ape cohabitation and differential human perceptions of gorillas and chimpanzees, a greater willingness to hunt gorillas, gorillas’ higher relative density and greater propensity to raid forest gardens. Interpreted together, our analyses in Cameroon point to two mechanisms facilitating such viral sharing: first, physical contact through great ape hunting, meat handling, and fecal contact, and second, environmental contact via focused co-use of small forest gardens by humans and gorillas.

## MATERIAL AND METHODS

### Study sites and study periods

We conducted research in several villages located between Yokadouma and Moloundou, Cameroon, and at the European zoo. Details of these two study sites are available elsewhere^22^. Team researchers (Victor Narat, Stephanie Rupp, Philippe Ambata, and Tamara Giles-Vernick) made a total of four field trips to Cameroon in 2015, 2016 and 2017 for six months total. We collected data and samples in the European zoo during a five-day research visit in November 2017. To protect the anonymity of human populations and activities, we do not report the names of individual study sites.

### Data collection

#### Qualitative data collection

We collected qualitative data using anthropological participant-observation and semi-directed interviews (Supplementary method 1) with inhabitants of forest villages (Cameroon) and zookeepers (European zoo). In Southeastern Cameroon, we collected 93 in-depth individual and collective interviews with 83 men and 31 women, complemented by many informal discussions and 150 hours of participant-observation of forest activities and recitation of mythical tales.

Participant-observations enabled us to observe human interactions with and in proximity to chimpanzees and gorillas. Interviews permitted us to collect qualitative data concerning gorilla and chimpanzee behaviors, diet, habitats, and interspecies contacts, as well as human perceptions and practices with these animals. Participant-observation and semi-structured interviews were conducted in French or in the Bangando language; all interviews were recorded. We collected detailed notes for participant-observations.

In the zoo, Victor Narat and Tamara Giles-Vernick observed and documented zookeeper-great ape interactions and living conditions among the great apes sampled, including feeding regimens, living conditions, and cleaning practices of their habitats. We also observed and conducted semi-structured interviews with zookeepers to document great ape contacts with other animal species, including humans.

#### Participatory longitudinal survey and transects in Cameroon

We developed a longitudinal participatory survey for Cameroon volunteers and conducted this survey over 10 months. Eighteen volunteers (8 women and 10 men over 21 years old) collected daily data on their contacts with gorillas and chimpanzees. Although a prior study addressed human-NHP physical contact^41^, the present article uses physical, direct and environmental contact data for gorillas and chimpanzees only (Table 1).

#### Questionnaire

Drawing from preliminary analyses of our qualitative data and participatory longitudinal survey, we developed and conducted a questionnaire (449 participants in four villages) to assess human contact frequency with gorillas and chimpanzees. Participants were asked if within the last day, week, month, year, or more than one year, they had had specific types of physical and direct (seen alive, heard) contacts with gorillas or chimpanzees. They also were asked to report on great ape involvement in crop raiding, namely the last great ape species raiding one of their fields, as well as the frequency of field raiding.

#### Sample collection

In Cameroon, after identifying great ape nesting sites, we collected fresh stool samples during the early morning, taking one stool per nest for a total of 15 gorilla samples and 6 chimpanzee samples. At the zoo, we worked with zookeepers to collect stool samples immediately after feces emission, for a total of 6 samples for each species.

For human stool collection in Cameroon, we included adult participants over 21 years old with no current or chronic health problems. All participations received an information notice and informed consent form, explained orally in the Bangando language. Among potential participants, we conducted a questionnaire to determine whether they suffered from a recent or chronic illness or used medicines (including antibiotics), and what forest-based activities they pursued. All participants received a sampling tube which they used for individual stool collection. All stool samples were delivered within 12 hours of emission and were subsequently stored in RNAlater^®^ tubes (Thermo Fisher)^22^. We successfully collected 24 human samples.

For human stool collection at the European zoo, we invited zookeepers working full-or part-time with gorillas or chimpanzees to participate in this study. Stool collection followed the same procedure as in Cameroon, although zookeeper participants stored the sample directly in the RNAlater tube and sent immediately in a secured package the Microbiology Service, Saint Louis Hospital in Paris, France.

All stool samples were frozen at −80°C within the 10 days of collection (for Cameroonian samples at Centre Pasteur, Yaoundé, and for zoo samples at Saint Louis Hospital, Paris). Further details of stool collection can be found in a previous publication^22^.

A total of 64 stools samples were analyzed, including 30 from humans (23 from Cameroon forest, 7 from European zoo), 21 from gorillas (15 from Cameroon forest, 6 from zoo), and 12 from chimpanzees (6 from Cameroon forest, 6 from zoo). Two samples, one from a Cameroon human (n°38) and another from a zoo chimpanzee (n°61), were removed from final analysis because the sequencing depth was too low (number of total reads < 5 million). We recovered a median of 40.6 million of reads per sample (range: 4.11E+06 – 5.79E+07) for DNA sequencing and of 40.7 million of reads per sample (range: 1.58E+06 −1.44E+08) for RNA sequencing. Samples and sequencing information are shown in Supplementary Table 1. The median read per million (RPM) of viral reads per sample was 536 (range 38 – 97251, IQR: 1372–367) (Supplementary Table 2).

### Data analyses

#### Contact frequency with great apes in Cameroon

We obtained daily contact data for 18 volunteers over 288 days on average (+/− SD=21.4, 230-303). We used presence/absence coding of data for one day, one volunteer, one type of contact and one great ape species. We then analyzed the frequency of contact (%) for each type of contacts, each species and each volunteer. We calculated the mean frequency for each type of contacts.

We calculated an estimated frequency of contact (%) using questionnaire data. We assigned a value that was the inverse of the number of days: 100 for daily contact, 14 for once weekly, 3 for once monthly, 0.3 for once yearly and 0.1 for more than once each year (estimated here at once every three years for coding). Based on this scale, we calculated the mean frequency for each type of contact.

For our longitudinal survey and questionnaire datasets, we compared contact frequencies with the two great ape species using a Mann-Whitney statistical test.

To calculate crop raiding frequency, we analyzed the proportion of respondents who identified a chimpanzee or gorilla as the raiding species for last time their field or garden was pillaged. We also calculated the proportion of respondents citing chimpanzees or gorillas as the species more frequently responsible for crop raiding.

#### Qualitative data analyses in both sites

Following transcription into French of all recorded interviews, we conducted manual coding of all qualitative data (transcriptions and notes), organizing data segments into categories pertaining to descriptions and perceptions of, discrete practices around, and interactions with great apes and other NHPs. From these codes, we used Thematic Analysis to identify broader, cross-cutting themes pertaining to human-great ape engagements.

#### DNA and RNA virome analysis with shotgun Next Generation Sequencing

Fecal samples (solid phase) were re-suspended and diluted (50%) in phosphate buffered saline (PBS) and then centrifuged at 2500 g for 20 minutes. The supernatant was then filtered using an 0.45μM filter. To enrich for viral particles by reduction of host background, stool supernatant was filtered through a 0.45-μm filter (Corning Costar Spin-X centrifuge tube filters), and an aliquot of 315 μl of filtrate was pretreated before extraction by incubation with different nucleases: TURBO DNase (Invitrogen, Carlsbad, CA); Baseline-ZERO DNase (Ambion, Foster City, CA); Benzonase (NEB); RNAse A (Promega) for 30 min, at 37 °C. Total nucleic acids were extracted using Nuclisens EAsymag (Biomerieux) according manufacturers protocol. For DNA libraries preparation, 25μL of extract was used. Depletion of methylated host DNA was performed using NEBNext^®^ Microbiome DNA Enrichment Kit (NEB) according to the manufacturer instructions. DNA was then purified using zymo DNA Clean (Zymo) and eluted in 7.5μL of sterile water. DNA libraries were prepared using Nextera XT library preparation kit (Illumina). For RNA libraries preparation, Trio RNA Kit (Nugen) was used according to manufacturer instructions. Libraries were sequenced on an Illumina HiSeq X using 150/150- bp paired-end sequencing.

#### Bioinformatics analysis

Raw reads were cleaned using TRIMMOMATIC^65^. Duplicated reads were removed using Dedupe^66^. Taxonomic assignment was carried out using Kraken2 with Viral, Bacterial and Human Refseq databases^67^. Kraken viral assigned reads were verified using Blastn on Refseq viral database. Reads with inconsistent assignment between Kraken and Blast methods were removed.

Alpha and Beta Diversity analysis were conducted using packages Phyloseq v1.22.3 and Vegan v2.5-4 in R v3.4.4^68,69^. For Alpha diversity, Simpson, Shannon, Chao1 and Richness index were calculated. For Beta diversity, Bray Curtis dissimilarity and Unifrac metrics were used. Principal coordinate analysis (PCoA) and network analysis was done with either Bray Curtis or weighted Unifrac distance. Permutational analysis of variance (PERMANOVA) was used to compare microbial communities between each group based on Bray Curtis dissimilarity indices, weighted and unweighted Unifrac distances using the adonis2 function with R package vegan.

Viral taxa shared between the human group and at least one great ape group (chimpanzee or gorilla) in Cameroon and in the zoo were identified.

As the genomic coverage of shared viruses between humans and apes did not allow the use of standard phylogeny methods, an alternative approach was used. Viral reads of interest were assembled into contigs with Spade^70^. Generated contigs and reference genomes (Supplementary Table 3) were compared using *All versus all BLAST*. Sequence similarity was assessed with the Bit-score and presented through sequence similarity networks with Cytoscape^71^.

## DATA AVAILABILITY

RNA and DNA Metagenomic raw data are deposited on Sequence Read Archive (pending accession number). All other data are available upon reasonable request, except for qualitative data which cannot be shared because of ethical restrictions.

## Acknowledgement

We are grateful to the inhabitants and authorities in southeastern Cameroon for their warm welcome and their support of this study. We also acknowledge the invaluable assistance of the European zoo and its employees where we conducted the investigation. We also greatly appreciate the contributions of Olivia Cheny of the Center for Translational Science at the Institut Pasteur. The French Agence Nationale de la Recherche (ANR-31-CE31-0004), CIFAR, the INCEPTION project (PIA/ANR-16-CONV-0005), and the National Endowment for the Humanities (US) provided funding for this study.

## Author contributions

VN contributed to study design, data collection, data encoding, data analyses and manuscript writing

MS contributed to study design, virome analyses and manuscript writing

MK and TH contributed to data encoding and analyses

SMD and NR contributed to virome analyses

SR contributed to study design, data collection and qualitative data analyses

PA contributed to data collection

RN contributed to study design and data collection

FS contributed to study design and virome analyses

JLG contributed to study design, virome analyses and manuscript writing

TGV contributed to study design, data collection, data analyses and manuscript writing and editing

## Competing interests

The authors declare to have no competing interests.

## Materials and correspondence

Tamara Giles-Vernick

Victor Narat

## Notes

### Competing Interest Statement

The authors have declared no competing interest.

