## Supplemental materials for "Unexpected higher convergence of human-great ape enteric viromes in central African forest than in a European zoo: A One Health analysis"

### Supplementary Table 1: Samples and sequencing information

Excel file “Supp_Table_1.xlsx”

### Supplementary Table 2: Viral reads information among the different groups

| | Min. | 1st Q | Median | Mean | 3rd Q | Max. |
|---|---|---|---|---|---|---|
| CamChim | 1077 | 1228 | 2027 | 17795 | 3074 | 97252 |
| CamGor | 42.56 | 103.40 | 358.80 | 495.59 | 636.78 | 2192.54 |
| CamHum | 38.33 | 70.01 | 232.17 | 2052.74 | 578.30 | 19386.06 |
| ZooHum | 95.58 | 150.85 | 265.97 | 8024.99 | 5812.55 | 43886.62 |
| ZooChim | 632.2 | 677.9 | 764.4 | 1226.1 | 1698.1 | 2357.7 |
| ZooGor | 3134 | 3648 | 16152 | 19561 | 30109 | 47685 |

All reads were exprimed in RPM (read per million of sequenced reads))

**Supplementary Table 3: Accession numbers of viral sequences used for the network analysis in Figure 4**

| Name | Accession Number |
|---|---|
| HAdV-A12 | NC001460.1 |
| HAdV-A18 | GU191019.1 |
| HAdV-A31 | AM749299.1 |
| HAdV-B11 | NC011202.1 |
| HAdV-B14 | AY803294.1 |
| HAdV-B16 | AY601636.1 |
| HAdV-B21 | AY601633.1 |
| HAdV-B3 | NC011203.1 |
| HAdV-B34 | AY737797.1 |
| HAdV-B35 | AY271307.1 |
| HAdV-B7 | AC000018.1 |
| HAdV-C1 | AF534906.1 |
| HAdV-C2 | NC001405.1 |
| HAdV-C5 | AC000008.1 |
| HAdV-C6 | HC492785.1 |
| HAdV-D36 | GQ384080.1 |
| HAdV-D37 | DQ900900.1 |
| HAdV-D46 | AY875648.1 |
| HAdV-D48 | EF153473.1 |
| HAdV-D49 | DQ393829.1 |
| HAdV-D53 | FJ169625.1 |
| HAdV-D54 | NC012959.1 |
| HAdV-D8 | AB448767.1 |
| HAdV-D9 | NC010956.1 |
| HAdV-E4 | NC003266.2 |
| HAdV-F40 | NC001454.1 |
| HAdV-F41 | DQ315364.2 |
| HAdV-G52 | DQ923122.2 |
| SAdV-A1 | NC006144.1 |
| SAdV-A48 | HQ241818.1 |
| SAdV-A6 | CQ982401.1 |
| SAdV-B21 | AC000010.1 |
| SAdV-C31 | FJ025904.1 |
| SAdV-E22 | AY530876.1 |
| SAdV-E23 | AY530877.1 |
| SAdV-E24 | AY530878.1 |
| SAdV-E25 | AC000011.1 |
| SAdV-G1 | NC006879.1 |

| | |
|---|---|
| SAdV-G7 | DQ792570.1 |
| Enterovirus A | AY421760.1 |
| Enterovirus B | NC_038307.1 |
| Enterovirus C | V01149.1 |
| Enterovirus D | AY426531.1 |
| Enterovirus E | D00214.1 |
| Enterovirus F | DQ092770.1 |
| Enterovirus G | AF363453.1 |
| Enterovirus H | AF326759.2 |
| Enterovirus J | AF326766.2 |
| Enterovirus K | KX156158.1 |
| Enterovirus L | KU587555.1 |
| Rhinovirus A | FJ445111.1 |
| Rhinovirus B | DQ473485.1 |
| Rhinovirus C | EF077279.1 |

### Supplementary Figure 1: Distribution of viral reads according to virus host

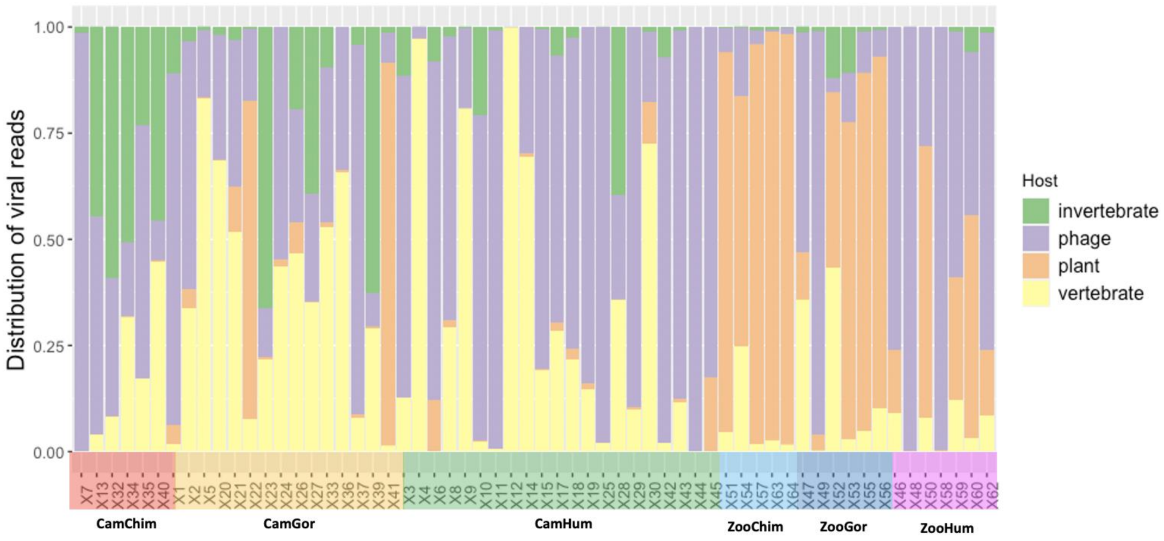

**Supplementary Figure 2: Boxplot of diversity indices for (a) all viruses and (b) vertebrate viruses.** ** indicates a p-value <0.05 (Wilcoxon test).*

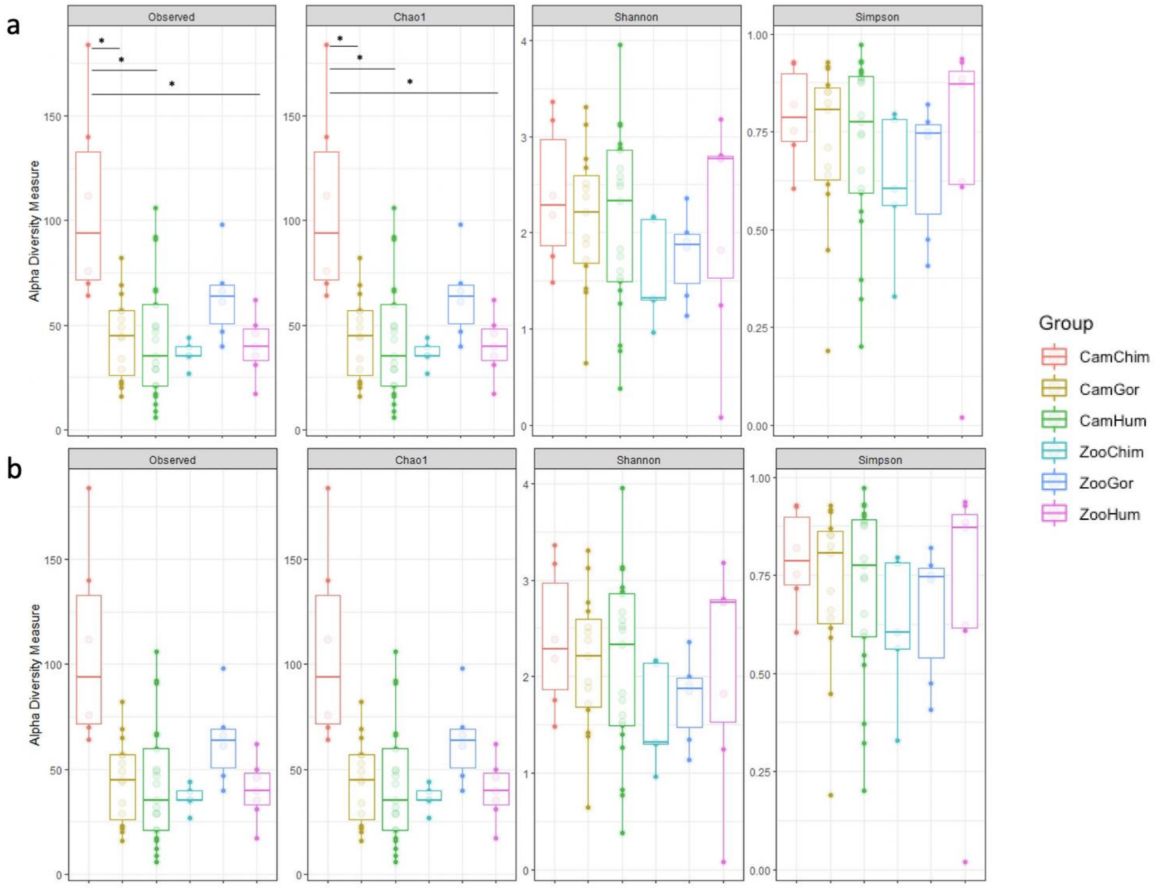

### Supplementary Methods 1: Semi-directed interview guides

For great ape zookeepers

1. How long have you worked at this zoo?
2. How long and in what capacity have you worked with this species of great ape?
3. Can you describe for me a typical day working with these animals?
4. Do your daily activities change seasonally, and if so, how?
5. Can you tell me more about feeding of these animals? (timing, dietary composition and quantities for each animal by age and sex, seasonal changes, other relevant practices)
6. From where do you obtain these foods?
7. How do you organize their time outdoors? Are there ever times when they do not go outdoors?
8. Do they eat plant material in the outdoor enclosure or drink from the canal around the island?
9. Do you provide additional supplements from time to time? What?
10. Can you describe any daily physical interactions that you have with these animals? Do you ever touch them? When and why?
11. Have these great apes been sick recently (within the last two months)? If so, which one(s)? What did you do? Did you provide treatment, and if so, what?
12. In your experience, is it possible for disease to pass between humans and captive animals in this zoo setting? Why or why not? Have you ever witnessed such transmission, and if so, when?

For inhabitants of southeastern Cameroon

1. What environmental changes have you observed or heard about in this forest? Over what period of time? What do you think of these changes?
2. In your language, how do you refer to a gorilla? A chimpanzee? Are there terms for referring to both gorillas and chimpanzees? Or terms to denote gorillas, chimpanzees, and monkeys?
3. Where do gorillas/chimpanzees reside in this forest? What are their preferred habitats? Do they remain in the same location, or do they move?
4. Do you find gorillas/chimpanzees in close proximity to your village (use map)?

5. Do you hunt gorillas or chimpanzees? Were these animals hunted in the past? When, and by whom?
6. Who hunts gorillas/chimpanzees now?
7. What weapons are used to hunt gorillas/chimpanzees?
8. How are gorillas or chimpanzees usually butchered? Where and by whom?
9. What do you do with the different parts of the animals?
10. Do people currently keep chimpanzees or gorillas as pets? Of what age? Why or why not? Has this practice of petkeeping changed over time?
11. Can you describe for me the qualities/personality traits of gorillas/chimpanzees?
12. How do they behave with one another? Have you observed their behavior? What features strike you as most important?
13. Do gorillas/chimpanzees have certain capacities or powers? Can you describe them? Have these capacities or powers changed over time?
14. Do gorillas/chimpanzees have knowledge? Of what?
15. What do people think about these animals?
16. How do you behave when you meet one? Why?
17. How do gorillas/chimpanzees behave when they see you?
18. Do people and gorillas/chimpanzees share certain capacities or knowledge?
19. Do you think that relations between people and gorillas/chimpanzees have changed? If so, since when? And why?
20. Do gorillas/chimpanzees ever fall ill? How do you know that they are ill? What are their symptoms?
21. What do gorillas/chimpanzees do when they are ill? How do they care for themselves? (If animal uses leaves for self-treatment, ask person to show leaves. Ask if people use these leaves as well.
22. Is it possible for an illness to be transmitted from a gorilla/chimpanzee to a human? Or from a human to a gorilla/chimpanzee? If so, which illnesses? How does this transmission take place?
